# Duplex Sequencing Provides Detailed Characterization of Mutation Frequencies and Spectra in the Bone Marrow of MutaMouse Males Exposed to Procarbazine Hydrochloride

**DOI:** 10.1101/2023.02.23.529719

**Authors:** Annette E. Dodge, Danielle P.M. LeBlanc, Gu Zhou, Andrew Williams, Matthew J. Meier, Phu Van, Fang Yin Lo, Charles C. Valentine, Jesse J. Salk, Carole L. Yauk, Francesco Marchetti

## Abstract

Mutagenicity testing is an essential component of health safety assessment. Duplex Sequencing (DS), an emerging high-accuracy DNA sequencing technology, may provide substantial advantages over conventional mutagenicity assays. DS could be used to eliminate reliance on standalone reporter assays and provide mechanistic information alongside mutation frequency (MF) data. However, the performance of DS must be thoroughly assessed before it can be routinely implemented for standard testing. We used DS to study spontaneous and procarbazine (PRC)-induced mutations in the bone marrow (BM) of MutaMouse males across a panel of 20 diverse genomic targets. Mice were exposed to 0, 6.25, 12.5, or 25 mg/kg-bw/day for 28 days by oral gavage and BM sampled 42 days post-exposure. Results were compared with those obtained using the conventional *lacZ* viral plaque assay on the same samples. DS detected significant increases in mutation frequencies and changes to mutation spectra at all PRC doses. Low intra-group variability within DS samples allowed for detection of increases at lower doses than the *lacZ* assay. While the *lacZ* assay initially yielded a higher fold-change in mutant frequency than DS, inclusion of clonal mutations in DS mutation frequencies reduced this discrepancy. Power analyses suggested that three animals per dose group and 500 million duplex base pairs per sample is sufficient to detect a 1.5-fold increase in mutations with >80% power. Overall, we demonstrate several advantages of DS over classical mutagenicity assays and provide data to support efforts to identify optimal study designs for the application of DS as a regulatory test.

## INTRODUCTION

Somatic mutations are a hallmark of cancer and exposure to mutagenic substances is associated with an increased risk of carcinogenesis (Jackson et al. 2018). Thus, testing the ability of chemicals to induce genotoxic effects is required prior to regulatory approval and a battery of internationally standardized assays is available for this purpose. There is also growing recognition that mutations in themselves represent an adverse effect that should be considered in regulatory decision-making (Heflich et al. 2020). Assays with available Organisation for Economic Co-operation Development (OECD) test guidelines are routinely employed as the gold standards to assess in vitro and in vivo mutagenicity (OECD 1997; 2022a; 2022b).

*In vivo* mutagenesis assays, such as the transgenic rodent (TGR) gene mutation assay (OECD 2022b), provide an important early indication of oncogenic risk; these assays are used to screen for potential hazard prior to the more time-intensive and costly rodent cancer bioassays. However, these assays have significant limitations. First, the TGR assay uses unique strains of animals that prevents integration with other standard toxicity tests. Second, the TGR plaque-based assay measures mutation induction in a single exogenous reporter gene that does not accurately represent the diverse structure and function of the mammalian genome. Many genomic features are thought to modulate differential sensitivity to chemical mutagens (Hodgkinson and Eyre-Walker 2011; Makova and Hardison 2015); these cannot be assessed when relying on a single reporter (Salk and Kennedy 2020). Third, the TGR assay, like many other existing mutagenicity tests, yields little mechanistic data. The pattern of mutation types induced by a chemical can provide clues to the mechanism behind the chemical’s mutagenic effect; with the TGR assay this is impractical given the need to laboriously pick and independently sequence many viral plaques (Beal et al 2020). Novel methodologies that can interrogate a broad selection of genomic targets and concurrently enumerate the spectrum of mutations across a diversity of genomic regions would provide mechanistic understanding of a chemical’s mutagenic effect for better prediction of human cancer risk.

Duplex Sequencing (DS) is an error-corrected next-generation sequencing (ecNGS) technology that detects ultra-rare mutations using a customizable panel targeting a portion of the endogenous genome (Kennedy et al. 2014). Recent work has demonstrated that DS is effective at measuring increases in mutations frequencies (MF) in mouse somatic tissues following exposure to established environmental mutagens including benzo[a]pyrene (BaP) (LeBlanc et al. 2022), aflatoxin (Chawanthayatham et al. 2017), N-ethyl-N-nitrosourea (ENU) and urethane (Valentine et al. 2020). In addition to being able to precisely quantify spontaneous and induced MF across diverse genomic contexts, chromatin states and functional domains, DS provides complementary mechanistic information in the form of detailed mutation spectral data (Salk and Kennedy 2020) that can be linked to mechanism-associated mutational signatures extracted from genome sequences of human cancers (Alexandrov et al. 2020) or cloned *ex vivo* treated cells (Kucab et al. 2019). For instance, BaP, the main carcinogenic agent found in cigarette smoke, produces a mutation spectrum that is highly similar to mutational signatures observed in human lung cancers (Nik-Zainal et al 2016; Kucab et al 2019; LeBlanc et al 2022). These signature analyses provide empirical evidence for the contribution of environmental mutagens to the mutation spectrum observed in human cancers. Given these features, it is conceivable that ecNGS may contribute to reducing reliance on, or even eliminating the need for, TGR and other mutation reporter assays.

Before DS can be used as a regulatory test for chemical mutagenicity assessment, more work is needed to compare its performance to standard assays across a range of chemicals with differing modes of action and potency. Here, we test the performance of DS compared to the TGR assay following exposure of mice to procarbazine hydrochloride (PRC), an anti-neoplastic agent that is classified as a Class 2A probable carcinogen by the International Agency for Research on Cancer (IARC Working Group 1981). PRC is well characterized as a potent *in vivo* clastogen; however, data on its mutagenic activity are limited (Maurice et al. 2019; Revollo et al. 2017; Suzuki et al. 1999). We measured mutations in the bone marrow (BM) of MutaMouse males exposed to PRC using the DS mouse mutagenesis panel comprised of twenty 2.4 kb targeted regions (Mouse Mutagenesis Kit, TwinStrand Biosciences Inc., Seattle WA, USA), chosen as a balanced representation of the entire genome with respect to location, GC content, trinucleotide abundances and genic status (LeBlanc et al. 2022). Our overall aim is to demonstrate the potential of DS as a robust *in vivo* mutagenicity test that could replace the TGR assay as the next “gold standard” in research and regulatory applications.

The objectives of this study are to: 1) use DS to study the dose-response of mutation induction by PRC in MutaMouse BM; 2) characterize regional variability in spontaneous and PRC-induced MF across the panel of 20 DS loci; 3) assess the concordance of DS with the *lacZ* TGR assay; 4) compare the mutation spectrum measured by DS to that measured by sequencing mutant *lacZ* plaques; and 5) explore optimal DS study designs for assessing mutagenicity by modeling sample size and sequencing data yield.

## METHODS

### Animal Exposure and Tissue Collection

All animal handling, exposures and tissue collection were approved by the Health Canada Ottawa Animal Care Committee and followed the Canadian Council on Animal Care guidelines. All MutaMouse animals used in this study were a random subset of those included in a previous TGR study investigating the effects of sampling time on mutant frequencies (Marchetti et al. 2021). Adult MutaMouse males, 8-10 weeks of age at the beginning of the exposure were exposed by oral gavage to 6.25, 12.5, or 25 mg/kg-bw/day PRC (CAS 366-70-1; Sigma-Aldrich, Oakville, ON Canada) or phosphate-buffered saline without calcium and magnesium (PBS; Corning cellgro, Manassas, VA, USA) as vehicle control (VC) for 28 consecutive days. Six mice for each treatment group, and three mice for VC were euthanized 42 days after the final dose. Three control mice sampled 70 days after the final dosing were added to achieve a matched control group of six animals. At euthanasia, BM was collected from the femurs, flash frozen, and stored at -80°C. BM was selected as the tissue to study because it is commonly used in regulatory mutagenicity testing and is a known target for PRC (Marchetti et al. 2021; Souliotis et al. 1994).

### Transgenic Rodent Gene Mutation Assay (lacZ assay)

The *lacZ* TGR assay was performed as per the OECD TG 488 (2022b). Genomic DNA was isolated from one femur using phenol/chloroform extraction (Gingerich et al. 2014). The quantity and purity of DNA were assessed using a NanoDrop spectrophotometer (Thermo Scientific Canada, Ottawa, Canada). λgt10 phage vectors in genomic DNA were excised using commercial packaging extract kits (Agilent Technology, Santa Clara, CA, USA), according to the manufacturer’s instructions. Transgene mutant frequency was determined using the phenyl β-d-galactopyranoside (P-gal)–positive selection assay (Lambert et al. 2005). A minimum number of 125,000 plaque-forming units (pfu) were counted per sample. Mutant frequency (phenotypically selectable mutants per *lacZ* locus) was expressed as the ratio of mutant plaques to total pfu.

### DNA Extraction and Library Preparation for Duplex Sequencing

DNA from the second femur was extracted using the Qiagen DNeasy blood and tissue kit, as described in the Qiagen User Manual (Cat. # 69504, Qiagen, Hilden, Germany). DNA quality was assessed using an Agilent Tapestation to ensure a DNA Integrity Number greater than 7.0. DNA was prepared for sequencing as described previously (Valentine et al. 2020) by TwinStrand Biosciences (Seattle, WA, USA). Briefly, 500 ng of DNA was ultrasonically sheared to a mean fragment size of approximately 300 base pairs (bp). The DNA was then end-polished, A-tailed and ligated to DuplexSeq™ Adapters (TwinStrand Biosciences, Seattle WA, USA).

The DS Mouse Mutagenesis panel has one target on each autosome, except for chromosome 1 that has two targets. Nine of the targets are located within genic regions and 11 within intergenic regions (Suppl. Table 1). DS targets were classified as genic or intergenic based on gene intersection information obtained from the gene database GenCode, the Mouse GenCode gene set, version M25 (Frankish et al. 2019). The chosen targets are presumed to be selectively neutral, with none of the targeted regions having known roles in cancer either in mouse or human orthologs. The panel was chosen for optimal performance in hybrid capture and contains no significant pseudogenes or repetitive elements that could potentially confound alignment or variant calling. Target chromatin status and %GC content were determined using the University of California Santa Cruz (UCSC) Mouse Genome Browser, reference genome mm10 (Kent et al. 2002). Descriptions of the 20 targets are provided in Supp. Table 1. The panel was captured using 120mer biotinylated oligo probes (Integrated DNA Technologies, Coalville, IA, USA).

**Table 1.** Mean mutant frequency estimates and corresponding pairwise comparisons between dose groups for the *lacZ* assay obtained using a GLM.

| **Dose (mg/kg-bw/day)** | **Mutant frequency (x 10<sup>-5</sup>)** | **SE (x 10<sup>-5</sup>)** |
| --- | --- | --- |
| 0 | 7.3 | 1.76 |
| 6.25 | 12.7 | 2.48 |
| 12.5 | 24.9 | 3.07 |
| 25 | 46.7 | 4.14 |
| **Pairwise comparisons** | **Fold Changes (± SE)** | **p-value\*** |
| 6.25 to VC | 1.74 ± 0.54 | 8.84 x 10<sup>-2</sup> |
| 12.5 to VC | 3.39 ± 0.91 | **4.05 x 10<sup>-4</sup>** |
| 25 to VC | 6.37 ± 1.63 | **1.54 x 10<sup>-6</sup>** |

\*bold font indicates p-value < 0.05

Prepared libraries were sequenced on a NovaSeq 6000 (Illumina, Sand Diego CA, USA) to a mean on-target duplex depth of ∼16,000x. The resulting sequence data were processed through the DS pipeline (TwinStrand Biosciences) as described in Valentine et al. (2020). The datasets used in the current study are available in the Sequence Read Archives (SRA) <u>BioProject ID PRJNA909196</u>. Bioinformatic processing involved extracting Duplex Tags, read alignment, grouping reads by their unique molecular identifier, quality trimming, error-correction of read groups by duplex consensus calling, consensus post-processing, re-alignment, and variant calling. Variants with a variant allele fraction >0.01 were considered germline mutations and were filtered out. Resulting data from the DS pipeline included a summary of sequencing quality metrics, mutation frequency (MF), simple mutation spectra, trinucleotide frequency, and MF per each 2.4 kb target.

Mutations that occurred more than once in a single sample were considered under two opposing assumptions: 1) all identical mutations represent the clonal expansion of a single mutational event; or 2) all identical mutations occurred as independent mutational events, possibly due to local genomic features that render it more susceptible to mutation. Making the former assumption and counting multi-read mutations as a single mutation in the MF calculation guards against over-estimating mutant frequency that could otherwise be artificially increased by clonal expansion. This calculation is the *minimum* mutation frequency (MF_Min_) that could explain the data. Alternatively, counting all identical mutations independently guards against the possibility of undercounting true independent mutations. This calculation is the *maximum* mutation frequency (MF_Max_) that could explain the observed data. Since it is rarely possible to be certain which of these two assumptions apply to any given mutation (and it could be a mixture of both), here, we consider both alternatives. It is also worth noting that clonal expansion of adverse mutations can lead to genetic disease and thus such events warrant consideration.

### Mutation Data Interpretation and Statistical Analysis

First, mutation induction was measured as mutant frequencies (phenotypically scorable mutants per *lacZ* locus) for the TGR *lacZ* assay and as MF_Min_ or MF_Max_ (represented as mutations per bp) for DS using the entire mouse mutagenesis panel. Estimated frequencies by dose were obtained for both assays using generalized linear models (GLM) in the R software with the quasibinomial distribution. Pairwise comparisons were conducted using the doBy R library (Højsgaard and Halekoh 2018) and these estimates were then back-transformed. Here, the delta method was employed to approximate the back-transformed standard errors (standard error of the mean; SEM) of the estimated MF. A Holm-Sidak correction was applied to adjust p-values for multiple comparisons for both MF_Min_ and MF_Max_. Next, we estimated MFs for each 2.4 kb target using generalized linear mixed models (GLMM) with a binomial error distribution in the R software. Pairwise comparisons based on dose and location relative to genes were conducted using the doBY R library as described above. The MF_Min_ and MF_Max_ response for each target was then compared individually against the TGR lacZ assay.

The mutation spectra data generated by DS was evaluated by considering first only the mutated base and then by extending the analyses to include the flanking bases. To determine which mutation substitution types differed between control and treated groups, a modified contingency table approach was used as described by (Piegorsch and Bailer 1994). Pairwise comparisons as described above were used to determine which mutation types were significantly increased from VC. Subsequently, we generated a 96-trinucleotide mutation spectrum and used cosine similarity to determine the Catalogue of Somatic Mutations in Cancer (COSMIC) single base substitutions (SBS) signatures (Alexandrov et al. 2020) that most closely resembled the PRC mutation spectra, using the MutationalPatterns package in R (Manders et al. 2022).

### Benchmark dose comparison of LacZ and DS Results

Benchmark dose (BMD) modeling was conducted using the PROAST web application (version 70.1, https://rivm.nl/en/proast) to mathematically model the PRC dose-response curves for both assays. The BMD estimated in our study represents a dose that produces a 50% increase (i.e., the benchmark response; BMR) in *lacZ* mutant frequency, MF_Min_ or MF_Max_; this BMR was selected based on previous recommendations for genotoxicity assessment (White et al. 2020).

The BMD is a measure of the potency of a chemical, with a lower BMD indicating a more potent mutagen. To estimate the BMD, different mathematical models (Hill, exponential, reverse exponential, and lognormal) were applied to fit the dose-response data. The Akaike Information Criterion was used to select the model with the best fit. 95% confidence intervals based on model averaging were generated from 200 bootstrap runs and were used to statistically evaluate the differences between BMDs derived with DS versus the *lacZ* assay, and across genomic loci.

### Power Analysis on DS Study Design

We performed a power analysis to determine the minimum sample size and total number of duplex base pairs needed to observe a significant increase in MF using DS. First, we used a binomial distribution to randomly generate control samples based on parameters from our experiment. Hypothetical mutation counts were generated based on the observed average MF_Min_ from our VC group (1.31 × 10^−7^ mutations per bp) and the average total number of bases sequenced per animal (depth; 7.98 × 10^8^ bp). Over-dispersion was incorporated assuming a normal distribution with mean zero and standard deviation taken from the sample variance. A sample variance of 0.062 was estimated from the standard-deviation obtained from the GLMM used for per target dose-response analysis. We randomly generated a treatment group using the experimental parameters and a selected fold-increase. Next, we used a GLM to calculate the significance of the fold-change in MF between control and treated samples, as described above, and calculated the power of that analysis under the set conditions. Finally, the bisection method was used to estimate the minimum detectable effect size (MDES) of the simulated experiment yielding a power of 80% and a p < 0.05. We varied the sample size to calculate the power and MDES of the various conditions. Finally, we investigated the effect of sample variance and total duplex bases on our power results. We increased sample variance from 0.06 to 0.1 to roughly double our observed value and varied the total number of duplex bases. We then calculated the total number of duplex bases required to detect a 1.5-fold increase in MF with 80% power under each of the various simulated conditions.

## RESULTS

### LacZ Mutant Frequency

*LacZ* assay mutant frequency was measured by dividing the number of identified mutant plaques by the total number of plaques counted. On average, 14, 21, 51, and 100 mutants were counted for VC, 6.25, 12.5, and 25 mg/kg-bw/day PRC, respectively (Supp. Figure 1). PRC induced a significant increase in mutant frequency (× 10^−5^) relative to VC in the middle and high dose groups from 7.33 ± 1.76 mutations per *lacZ* locus (± SEM) in the VC to 24.9 ± 3.07 and 46.7 ± 4.14 at the middle and high PRC doses, respectively. No statistically significant increase was observed at the low PRC dose (Table 1). There was substantial inter-individual variation within the *lacZ* data, as indicated by the large SEM for the average mutant frequency of each dose group. These results are highly comparable to those reported by Marchetti et al. (2021) for the same animals.

**Figure 1.**
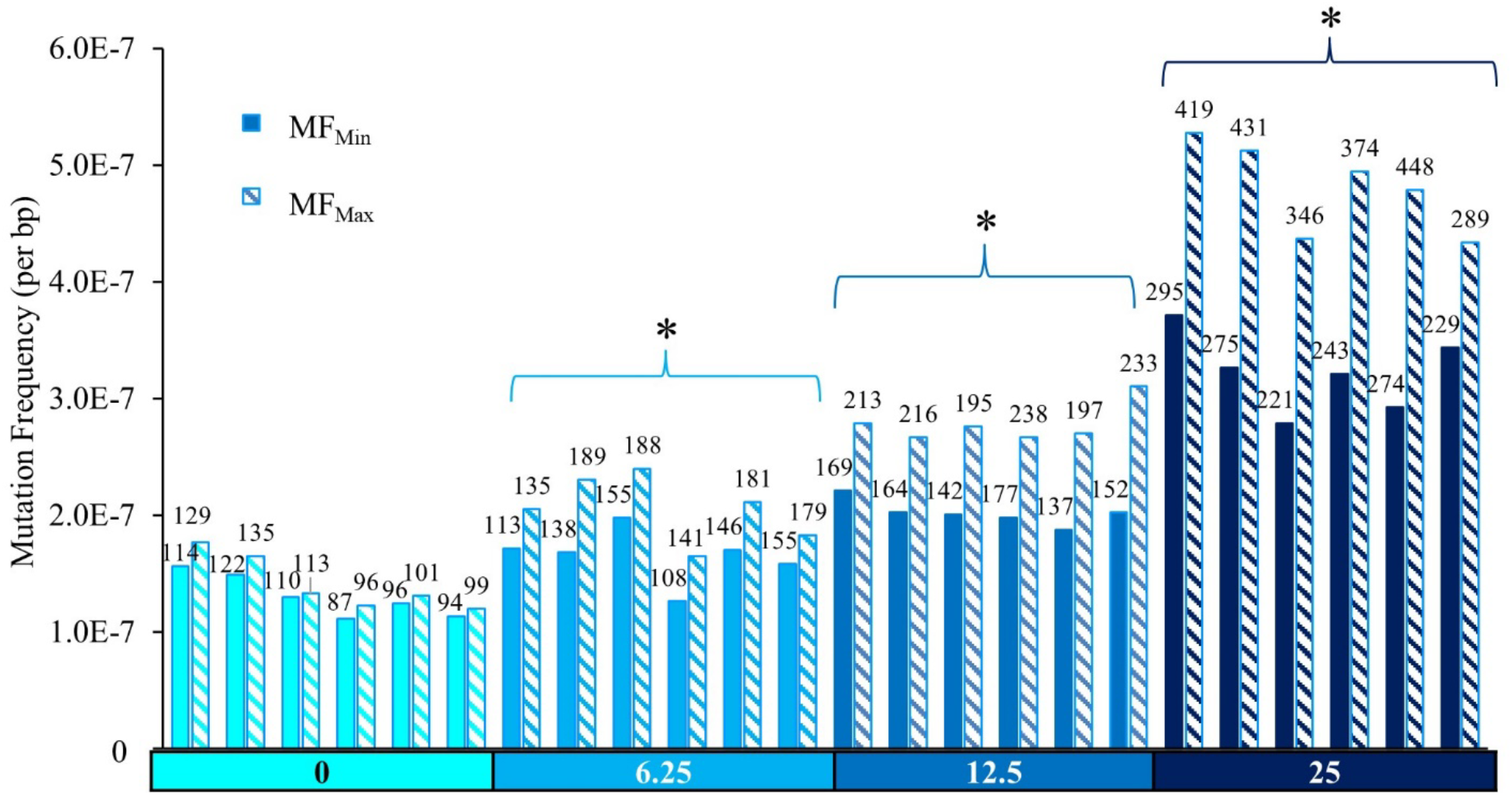
Duplex Sequencing mutation frequencies (MF) in the bone marrow of MutaMouse males at various doses of PRC. Bars represent the MF (mutations per bp) for each animal using MF_Min_ or MF_Max_. Data labels indicate total mutation count per animal. X-axis indicates PRC dose group (mg/kg-bw/day). Asterisks indicate a significant difference (p < 0.05 relative to the control in the average MF across animals).

### DS Mutation Frequency

DS reads were distributed relatively evenly across the 20 targets in the 24 samples (Supp. Figure 2). Sequencing yielded an average of approximately 800 million duplex base pairs per sample for a total of about 20 billion duplex bases across the cohort. All samples met a minimum target of 500 million duplex bases per sample.

**Figure 2.**
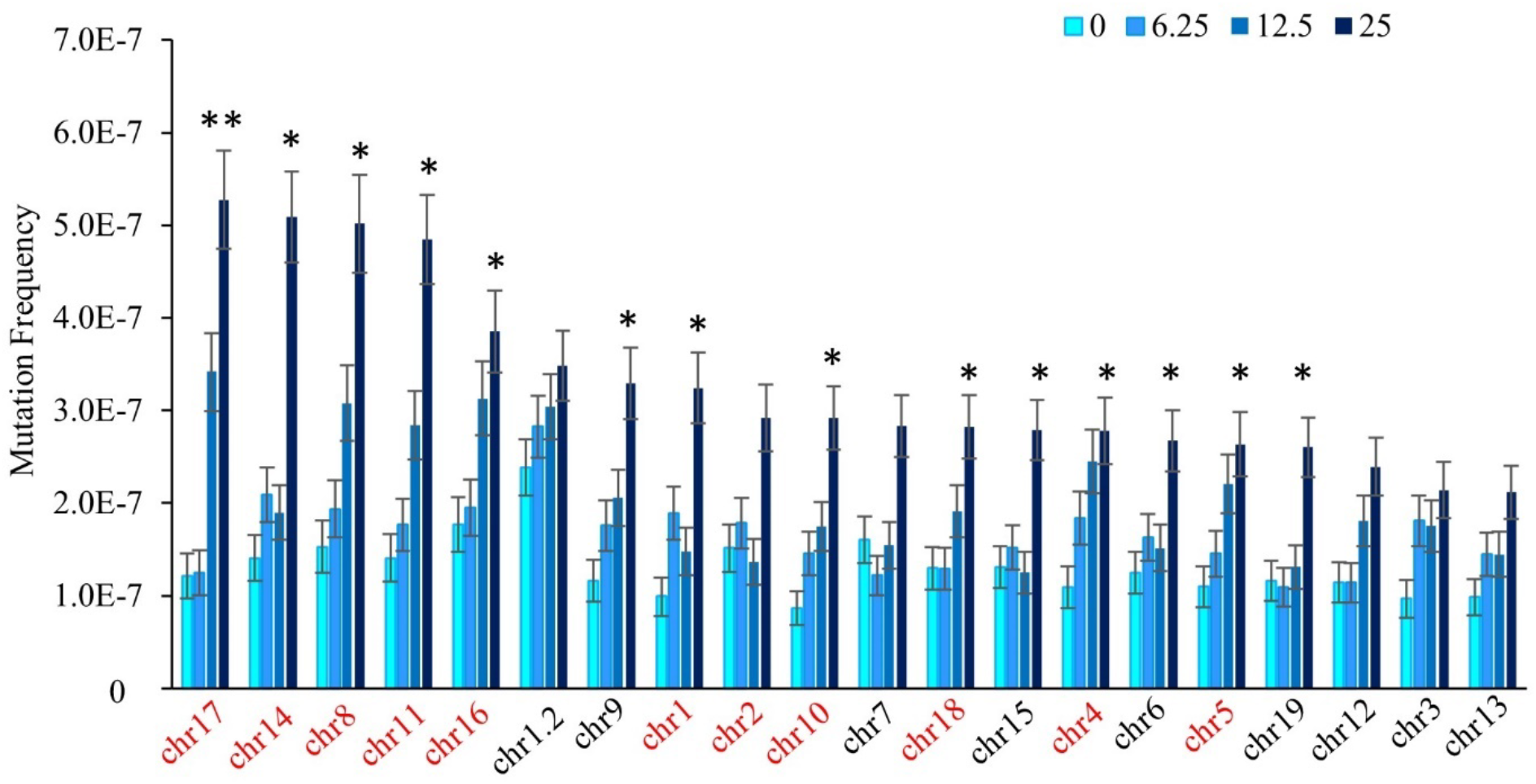
Spontaneous and PRC-induced MF by Duplex Sequencing target ordered from highest MF_Min_ in the high PRC dose group to the lowest. Data are mean MF ± SEM mutations per bp, for each target (n = 6), separated by dose (mg/kg-bw/day). Intergenic targets are labelled in red and genic targets are in black. Asterisks indicate a significant increase in the high PRC dose relative to the controls. The double asterisk indicates a significant increase in the high and middle PRC doses relative to controls. (Generalized linear mixed model, p < 0.05).

MF was calculated by dividing the number of identified mutants by the total number of bases sequenced per sample. Mutations include single nucleotide variants (SNVs), insertions and deletions (indels), and multi-nucleotide variants (MNVs). Under the conservative MF_Min_ assumption, 104, 136, 157, and 256 unique mutations on average were identified in mice treated with VC, 6.25, 12.5, and 25 mg/kg-bw/day PRC, respectively, with minimal inter-individual variability within each dose group (Figure 1). PRC induced a significant dose-dependent increase in MF_Min_ relative to VC in all PRC dose groups, from 1.31 × 10^−7^ ± 7.47 × 10^−9^ mutations per bp in the control to 3.22 × 10^−7^ ± 11.7 × 10^−9^ in the high dose group. A 1.3-, 1.5-, and a 2.5-fold increase in MF_Min_ relative to VC was induced in 6.25, 12.5, and 25 mg/kg-bw/day dose groups, respectively (Table 2).

**Table 2.**
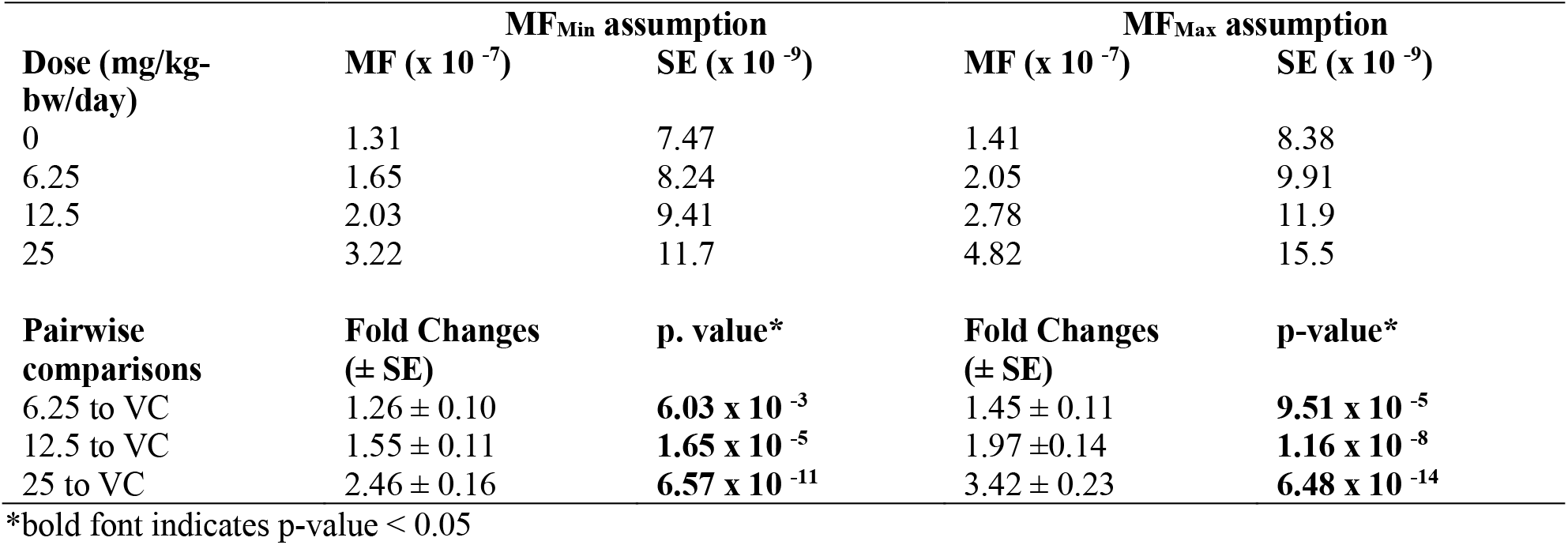
Mean mutation frequency (MF) estimates and corresponding pairwise comparisons between dose groups for DS obtained using a GLM.

| Dose (mg/kg-bw/day) | MF<sub>Min</sub> assumption |  | MF<sub>Max</sub> assumption |  |
| --- | --- | --- | --- | --- |
|  | MF (x 10<sup>-7</sup>) | SE (x 10<sup>-9</sup>) | MF (x 10<sup>-7</sup>) | SE (x 10<sup>-9</sup>) |
| 0 | 1.31 | 7.47 | 1.41 | 8.38 |
| 6.25 | 1.65 | 8.24 | 2.05 | 9.91 |
| 12.5 | 2.03 | 9.41 | 2.78 | 11.9 |
| 25 | 3.22 | 11.7 | 4.82 | 15.5 |
| Pairwise comparisons | Fold Changes (± SE) | p. value* | Fold Changes (± SE) | p-value* |
| 6.25 to VC | 1.26 ± 0.10 | **6.03 x 10<sup>-3</sup>** | 1.45 ± 0.11 | **9.51 x 10<sup>-5</sup>** |
| 12.5 to VC | 1.55 ± 0.11 | **1.65 x 10<sup>-5</sup>** | 1.97 ± 0.14 | **1.16 x 10<sup>-8</sup>** |
| 25 to VC | 2.46 ± 0.16 | **6.57 x 10<sup>-11</sup>** | 3.42 ± 0.23 | **6.48 x 10<sup>-14</sup>** |

\*bold font indicates p-value < 0.05

To assess the impact of counting all mutations independently, we performed the same analyses using the MF_Max_ assumption. We identified an additional 50, 198, 351, and 770 mutations for VC, 6.25, 12.5, and 25 mg/kg-bw/day PRC, respectively. PRC induced a significant dose-dependent increase in MF_Max_ (Table 2, p < 0.01) in all groups relative to VC from 1.41 × 10^−7^ ± 8.38 × 10^−9^ mutations per bp (± SEM) in the control to 4.82 × 10^−7^ ± 15.5 × 10^−9^ in the high dose group (Figure 1). A 1.5-, 2.0-, and a 3.4-fold increase in MF_Max_ relative to VC was induced in 6.25, 12.5, and 25 mg/kg-bw/day PRC dose groups, respectively. The MF_Max_ for the middle and high dose groups were significantly higher than to the MF_Min_ for those dose groups (p < 0.01). Inter-individual variability did not meaningfully differ between the MF_Min_ and MF_Max_ assumptions.

### DS Mutation Frequency by Target

Next, we analyzed mutation induction across the 20 genomic targets using the MF_Min_ assumption. The background MF_Min_ differed by 2.8-fold across the targets, with the highest MF_Min_ observed for the second target on chromosome 1 (chr1.2: 2.41 × 10^−7^) and the lowest MF_Min_ on chromosome 10 (0.858 × 10^−7^). PRC induced a significant increase in MF_Min_ relative to VC in 14 of the 20 targets at the high PRC dose (Figure 2). At the middle PRC dose, only the target on chromosome 17 had a significant increase in MF_Min_. No target had a significant increase in MF_Min_ compared to VC at the low dose. We observed a maximum 2.5-fold difference between the target with the highest MF_Min_ at the high PRC dose (chromosome 17; 5.21 × 10^−7^) and the target with the lowest MF_Min_ at the high dose (chromosome 13; 2.13 × 10^−7^). Similarly, we observed a 2.7- and 2.6-fold change between the targets with the highest and lowest MF_Min_ for the middle and low PRC doses, respectively. The targets with the highest MF_Min_ were on chromosome 17 for the high and middle PRC dose, and on the second target on chromosome 1 (i.e., chr1.2) for the VC and low PRC dose. Three of the five targets with the lowest background MF_Min_, those on chromosomes 3, 5 and 13, were also among the five targets with the lowest MF_Min_ at the high PRC dose, suggesting these targets may be less sensitive to mutation induction.

We observed that genomic targets in intergenic regions tended to have a higher susceptibility to PRC than targets in genic regions, which, if expressed, can be subjected to transcription-coupled repair (TCR). The five targets with the highest MF_Min_ at the high PRC dose were all located in intergenic regions, while 4/5 targets with the lowest MF_Min_ were genic (Figure 2). Overall, a combined analysis of intergenic targets showed a significantly higher mean MF_Min_ than the combined genic targets in the middle and high PRC dose groups by 1.3 and 1.4-fold respectively (p < 0.001) (Supp. Figure 3). These results did not change when using MF_Max_.

**Figure 3.**
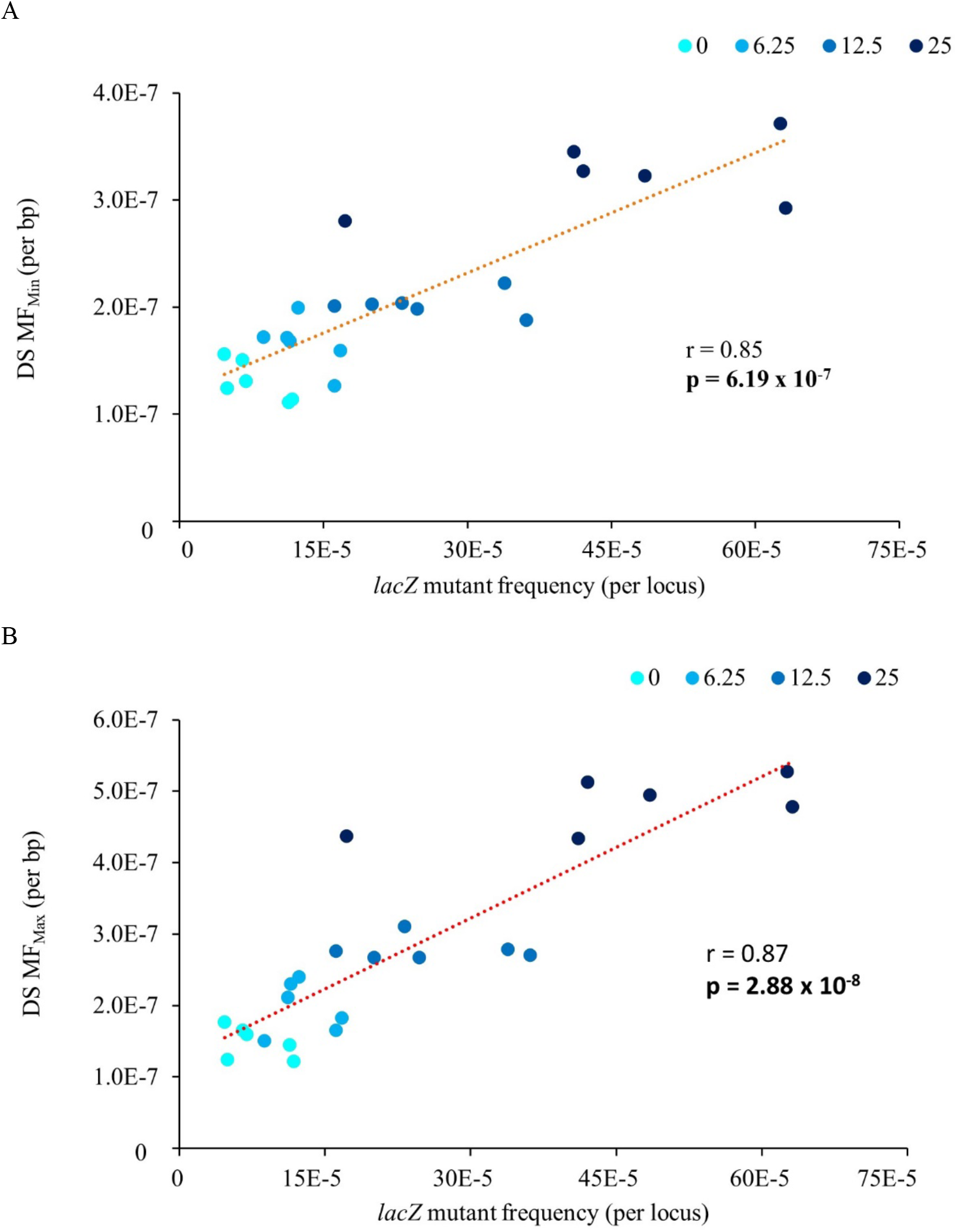
Pearson’s correlation analysis between *lacZ* mutant frequency and Duplex Sequencing using MF_Min_ (**A**) or MF_Max_ (**B**) for MutaMouse males exposed to PRC. Data are presented as the individual MFs for each animal.

Furthermore, intergenic targets showed a significant increase in MF_Min_ and MF_Max_ at all PRC doses compared to the VC, whereas the genic targets showed a significant increase only at the high dose for MF_Min_ and the middle and high dose for MF_Max_. Conversely, the mean background MF_Min_ observed in intergenic targets (1.26 × 10^−7^) was not significantly different from the mean background MF_Min_ in genic targets (1.28 × 10^−7^). These results were not changed when using MF_Max_.

### DS and TGR concordance

To assess DS concordance with the TGR assay, we compared the DS MF_Min_ to the *lacZ* mutant frequency in the same samples. There was a significant positive correlation between the induced DS MF_Min_ and *lacZ* mutant frequencies (Figure 3; Pearson’s correlation; r = 0.85, p = 6.19 × 10^−7^), with *lacZ* displaying more inter-individual variability. The correlation increased further when we used MF_Max_ (Pearson’s correlation; r = 0.87, p = 2.88 × 10^−8^), although the magnitude of the response to PRC was still higher in the *lacZ* assay (Table 1, Table 2). The *lacZ* fold changes relative to control for each dose group were greater by an average of 2-fold than DS MF_Min_ and 1.5-fold compared with DS MF_Max_.

We then assessed the concordance of MF of individual DS targets with the *lacZ* assay for the two PRC doses that significantly increased *lacZ* mutations. The fold-change in MF between VC and the PRC dose groups was estimated for each target, using either the MF_Min_ or MF_Max_ DS data (Figure 4). Using MF_Min_, the target on chromosome 17 (4.34 ± 0.98-fold-increase) was the only one within range of the fold-change observed using the *lacZ* assay (6.37 ± 1.63). Conversely, using MF_Max_, we observed six targets whose fold-increase in MF falls within the *lacZ* assay range, namely the targets on chromosome 17, 14, 1, 11, 8, and 10. These six targets were all intergenic. Similar trends were observed at the middle PRC dose, where four and 10 targets were within range of the *lacZ* fold-change (3.39 ± 0.91) when MF_Min_ and MF_Max_ values were used, respectively. Of these, all targets except for the one on chromosome 3 were intergenic (Supp. Figure 4). Thus, the data show intergenic targets generated fold changes in MF that more closely approximated *lacZ* assay results than genic targets.

**Figure 4.**
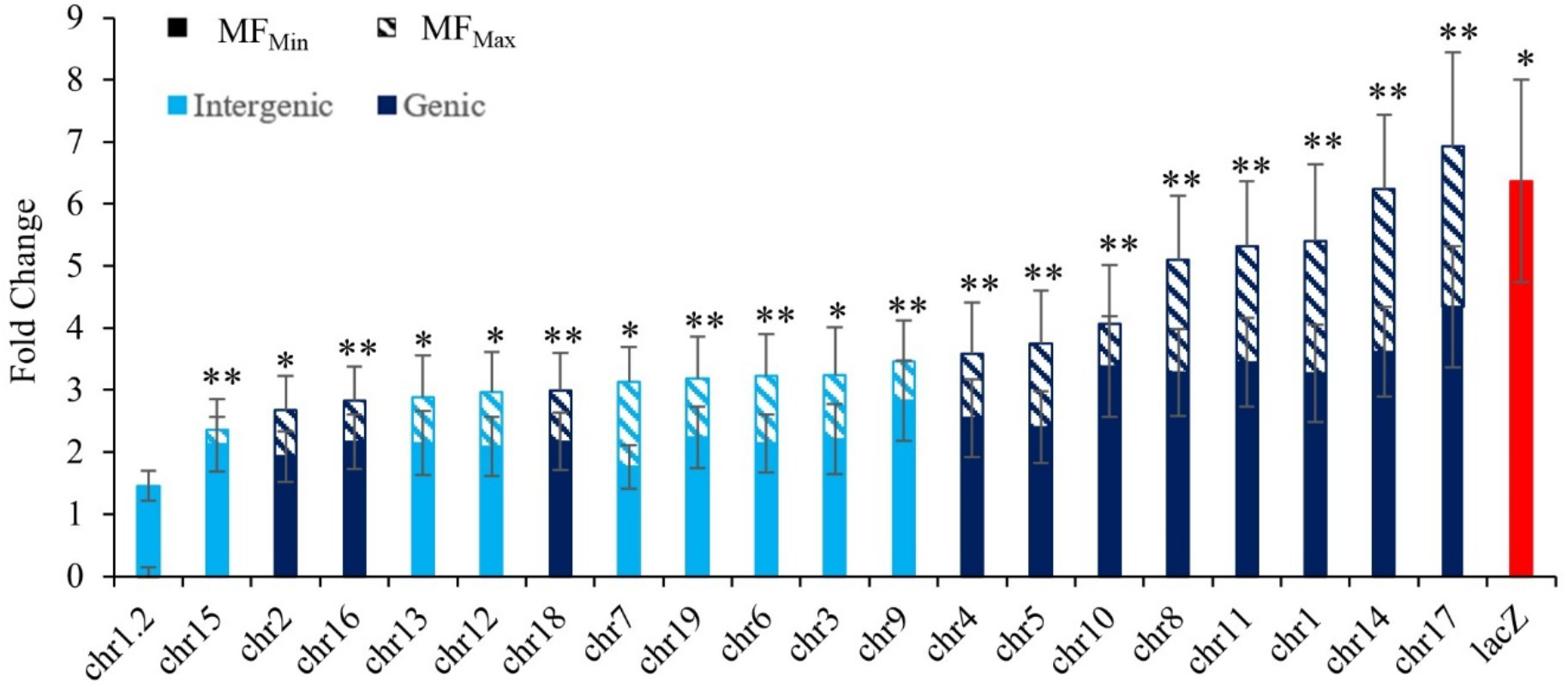
Fold change in MF between PRC high-dose group and VC for the 20 Duplex Sequencing targets as well as for the *lacZ* gene (red bar). Estimated using a general linear mixed model. DS targets are listed along the X-axis. Errors bars are SEM. Asterisks indicate the fold change was a significant increase in MF from VC for MF_Max_. Double asterisks indicate significance for both MF_Max_ and MF_Min_. The targets chr17, chr14, chr1, chr11, chr8, and chr10 were within range of the lacZ fold change.

Finally, we examined DS and *lacZ* concordance using BMD modelling to identify the dose at which a 50% increase in MF occurred above background. The BMD for the *lacZ* assay was 3.7 mg/kg-bw/day (95% CI of 1.6-6.6 mg/kg-bw/day), which was significantly lower than the BMD of 11.1 mg/kg-bw/day (95% CI of 8.7-13.8 mg/kg-bw/day) obtained using DS MF_Min_ (Figure 5A). However, using MF_Max_ produced a BMD of 8.4 mg/kg-bw/day (95% CI of 6.5-10.4 mg/kg-bw/day), which overlaps with the CI of the *lacZ* assay. Thus, quantitative concordance of DS with the *lacZ* assay improves when MF_Max_ are used. We also produced BMDs for each of the 20 DS targets individually calculated as MF_Max_ (Figure 5B) and compared these to the *lacZ* BMD. Every target had CIs that overlapped with the *lacZ* CI, except for targets on chromosomes 7, 15, and 19, which were significantly higher. Chr 1.2 failed to produce a BMD due to its weak dose-response and high mutation background frequency. Overall, BMD concordance between the assays improved when MF_Max_ was used.

**Figure 5.**
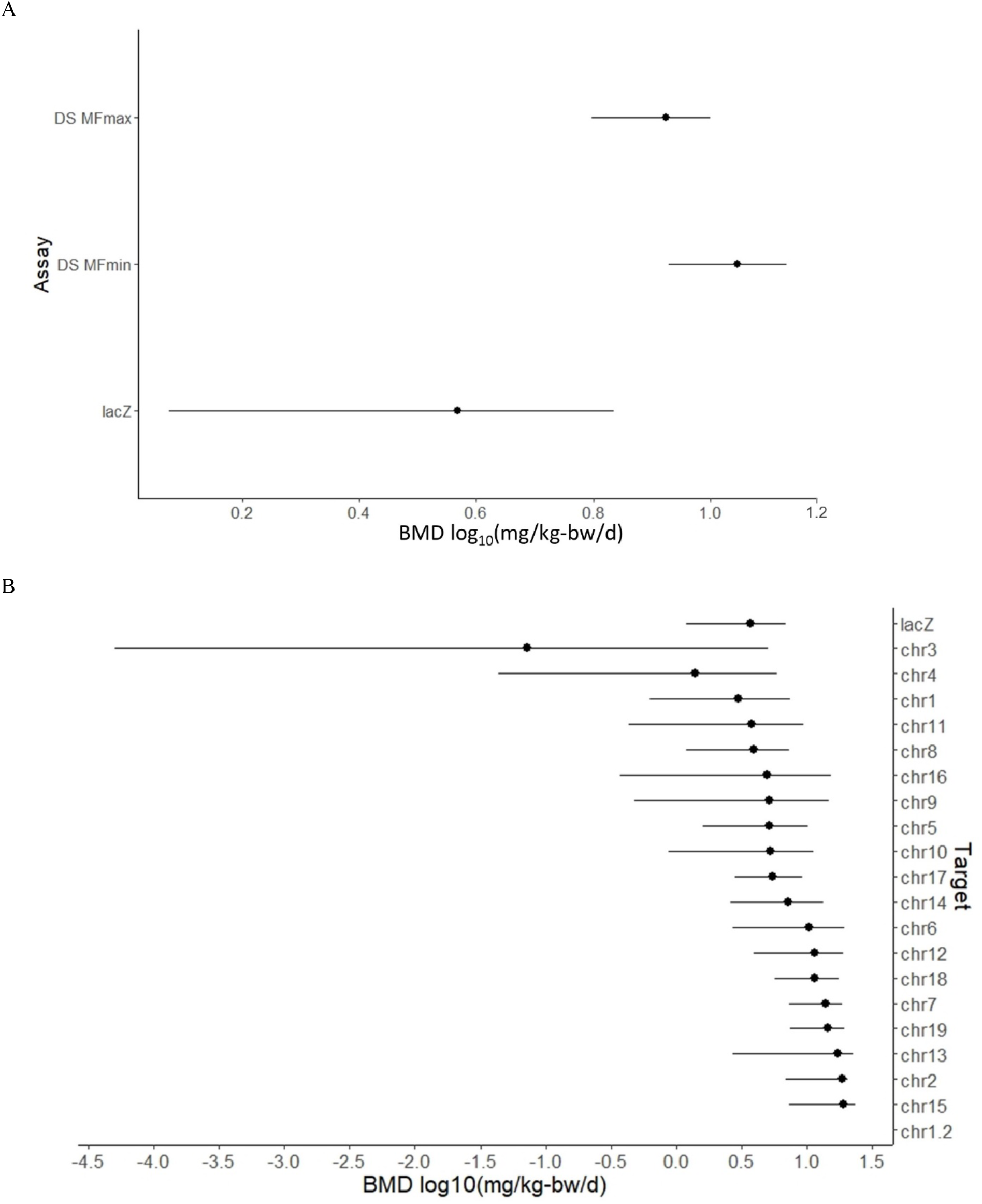
95% confidence intervals (CI) based on BMD model averaging analyses. (**A**) log_10_ BMD for a 50% increase of *lacZ* mutant frequency and the Duplex Sequencing MF, using either MF_Min_ or MF_Max_. (**B**) log_10_ BMD of *lacZ* mutant frequency and Duplex Sequencing MF_Max_ for each genomic target. DS targets are organized from smallest BMD to highest. Target chr1.2 failed to produce a BMD due to poor dose response.

### Mutation Spectrum

Both the spontaneous and PRC-induced mutation spectra (MF_Min_) consisted primarily of SNVs, followed by small deletions, then MNVs, and lastly, small insertions (Figure 6). Pairwise comparison of DS mutation data revealed that the overall mutation spectra were significantly different from each other at all PRC doses (modified contingency table; p < 0.05). PRC primarily induced T:A>C:G and C:G>T:A transitions accounting for 29% and 28% of total mutations at the high dose, followed by T:A>A:T transversions (13%) and T:A>G:C transversions (4%). This differed from the background mutation spectrum, which consisted primarily of C:G>G:C transversions (33%), C:G>A:T transversions (33%), and C:G>T:A transitions (17%). We observed significant dose-dependent increases in T:A>C:G transitions in all three dose groups compared with VC; T:A>A:T transversions were significantly increased at the middle and high PRC dose compared with VC; and T:A>G:C transversions and C:G>T:A transitions were increased at the high dose only (Figure 6).

**Figure 6.**
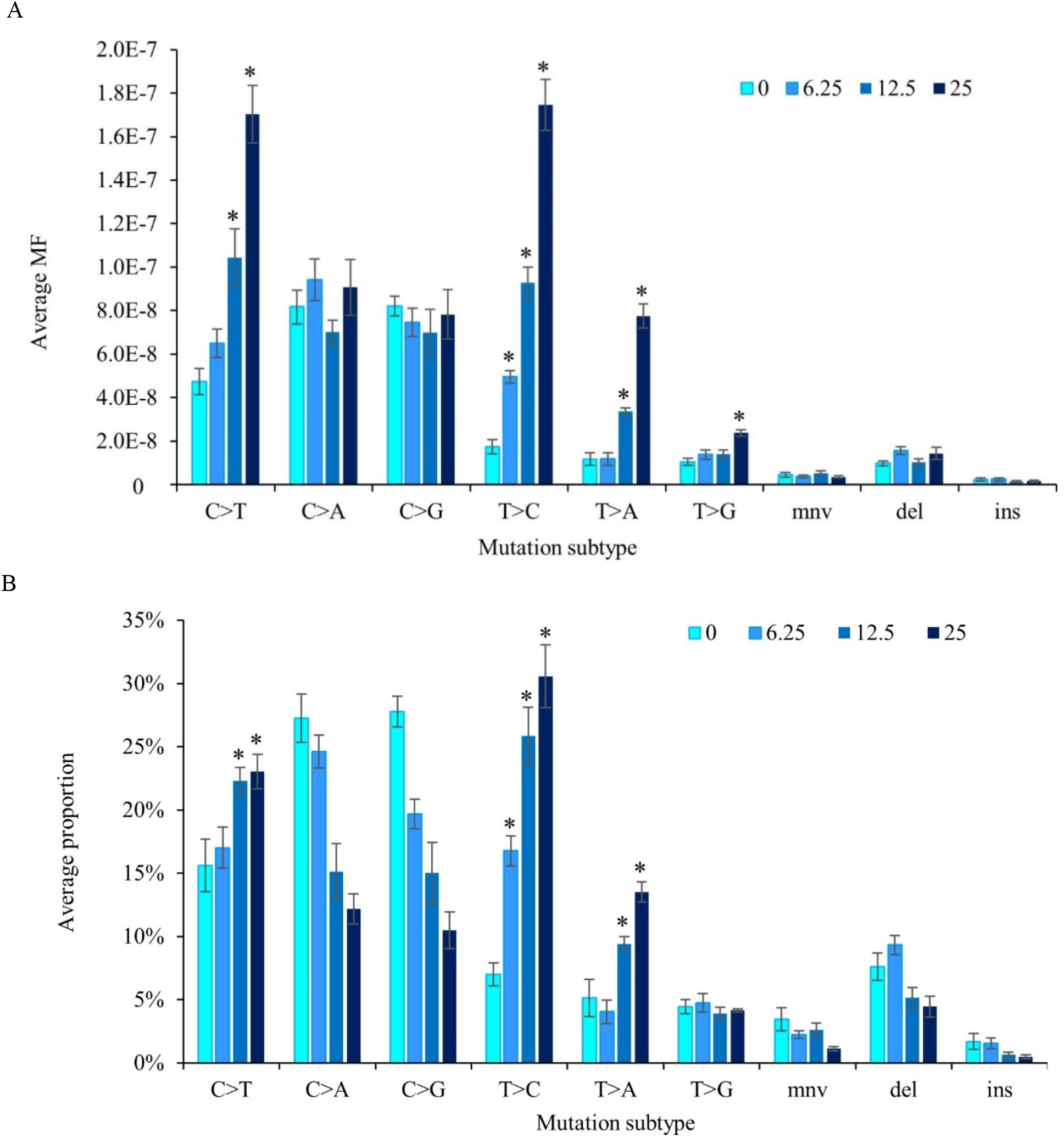
Mutation spectrum of controls and PRC dose groups measured by Duplex Sequencing in the bone marrow of MutaMouse males. (**A**) Mutation subtypes are presented as the average MF ± SEM. (**B**) Mutation subtypes are represented by the proportion of total mutations. Values are mean ± SEM. Plots represent MF_Min_. Mutation subtypes include the six single nucleotide variants using a pyrimidine reference, multi-nucleotide variants (mnv), small insertion (ins), and small deletion (del) mutations. Asterisks indicate a significant increase in MF from controls.

We further analyzed the sequence context of induced mutations by considering the flanking base on either side of the mutated base to produce trinucleotide spectra and examine mutational signatures (Alexandrov et al. 2013). This revealed a clear enrichment in T:A>C:T transitions and T:A>A:T transversions, together with a shift from mutations at CpG sites to non CpG sites for C:G>T:A transitions (Supp. Figure 5). We compared our trinucleotide frequencies to single-base substitution (SBS) signatures of the COSMIC Mutational Signatures database (mm10, version 3.3) using cosine similarity. We observed a dose-dependent increase in cosine similarity to several cancer signatures. The signature that displayed the greatest difference between its similarity to the control spectrum and its similarity to the high PRC spectrum was SBS 21, followed by SBS 26, and SBS 12 (Suppl. Figure 6).

### Power Analysis for Sample Size and Total Duplex Basepairs

We performed a power analysis on datasets simulated from the observed conditions of our DS experiment. Using our true sample size of six animals per dose, we had >95% power to detect a 1.5-fold increase in MF_Min_ (Table 3). Reducing the number to three animals per dose group retained 92% power. The MDES that could be achieved after reducing the sample size to three animals was a 1.4-fold increase in MF from background (Table 3). This is higher than the 1.26-fold increase in MF_Min_ that was observed for the lowest PRC dose, but is lower than the 1.55-fold and 2.46-fold increase observed at the middle and high PRC doses. Thus, under the observed conditions of our experiment, a reduction in sample size to three animals per group would result in sufficient power to classify PRC as mutagenic and detect the changes we observed at the middle (95% power) and high (>95% power) PRC dose but would be under-powered to detect the smaller increase observed at the low PRC dose (49% power). The use of MF_Max_ did not alter the results of the power analysis (data not shown). Finally, we assessed whether these results were robust to simulated increases in sample variance. We performed the same analyses as above but with twice the observed sample variance (0.1). We observed an 81% power to detect a 1.5-fold increase in MF_Min_ above background with a sample size of three and the increased sample variance.

**Table 3.** Power analysis for Duplex Sequencing dose-response studies.

| **Experimental conditions** |  |  |  |  | **Simulated condition** |  |
| --- | --- | --- | --- | --- | --- | --- |
|  | **MF<sub>Min</sub>** |  | **MF<sub>Max</sub>** |  | **Increased Variance** |  |
| **Background MF** | 1.31 x 10<sup>-7</sup> |  | 1.41 x 10<sup>-7</sup> |  | 1.31 x 10<sup>-7</sup> |  |
| **Total DS base pairs** | 7.98 x 10<sup>8</sup> |  | 7.98 x 10<sup>8</sup> |  | 7.98 x 10<sup>8</sup> |  |
| **Sample Variance** | 0.062 |  | 0.063 |  | 0.10 |  |
| **Sample Size** | **Power (%)** | **MDES<sup>b</sup>** | **Power (%)** | **MDES<sup>b</sup>** | **Power (%)** | **MDES<sup>b</sup>** |
|  | **1.5-FC<sup>a</sup>** |  | **1.5-FC<sup>a</sup>** |  | **1.5-FC<sup>a</sup>** |  |
| 6 | 100 | 1.23 | 100 | 1.22 | 100 | 1.27 |
| 5 | 100 | 1.25 | 100 | 1.25 | 98 | 1.31 |
| 4 | 100 | 1.30 | 99 | 1.29 | 95 | 1.38 |
| 3 | 92 | 1.40 | 91 | 1.38 | 81 | 1.51 |

<sup>a</sup> Power (%) to detect a 1.5-fold-change of MF<sup>b</sup> MDES: minimum detectable effect size with 80% power

Next, we investigated the number of total duplex bases required for analysis (Figure 7). When using a sample size of three animals, and our observed sample variance (0.062), a minimum of 500 million DS bases per animal is required to achieve sufficient power (81%) to detect a 1.5-fold increase in MF_Min_. However, this reduction in data yield causes analyses to become more susceptible to increases in sample variance. When the sample variance is 0.1, and the sample size is three animals, the power to detect a 1.5-fold increase in MF_Min_ is reduced to 71% and a minimum data yield of ∼800 million duplex bases is required to achieve sufficient power. Increasing the number of animals per group to four, with a minimum of 300 million duplex bases, would be sufficient to detect a 50% increase in MF_Min_ for datasets with a sample variance of 0.1 (78.2% power).

**Figure 7.**
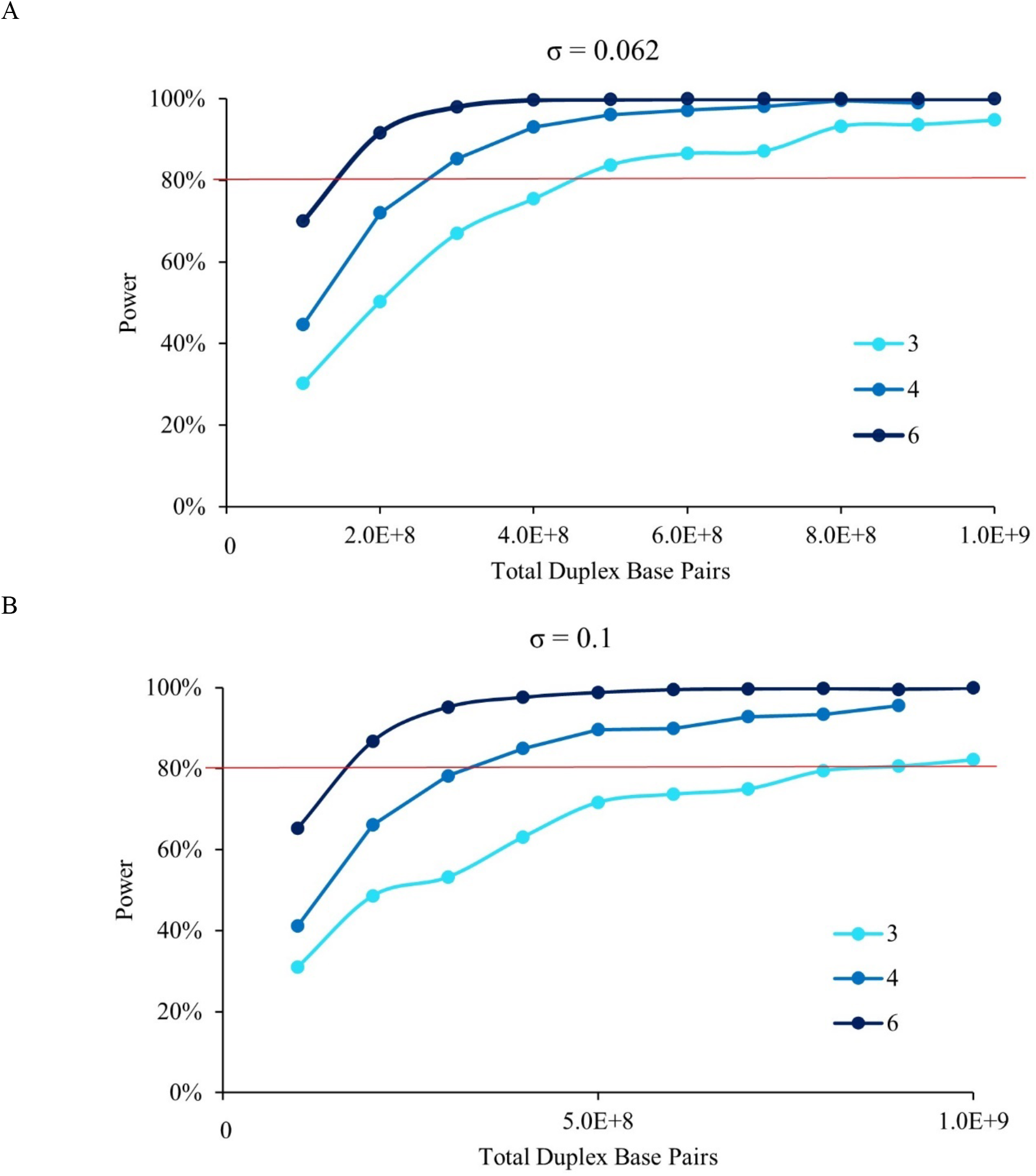
The power to detect a 50% increase in MF by Total Duplex Base Pairs. Results are displayed for a sample size of 6 animals per group (dark blue), 4 animals per group (blue) and 3 animals per group (light blue). (**A**) Sample variance of 0.062 (based on current study). (**B**) Sample variance of 0.1. Red line indicates the threshold for > 80% power and p < 0.05.

## DISCUSSION

We applied DS to study spontaneous and PRC-induced mutations in the BM of MutaMouse males across a panel of diverse genomic targets and compared the results with those obtained with the OECD TGR assay on the same samples. Low intra-group variability within DS samples resulted in statistically significant increases at all PCR doses, whereas the *lacZ* assay failed to detect a significant increase at the low dose. The *lacZ* assay initially showed a stronger fold-response to PRC than DS based on MF_Min_; however, when using MF_Max_, several DS targets had fold-increases consistent with the *lacZ* assay. DS revealed a PRC mutation spectrum that was distinct from the background spectrum, with PRC primarily inducing T:A>C:G transitions, followed by C:G>T:A transitions, and T:A>A:T transversions. Finally, power analyses suggested that reducing the sample size to three animals per dose group and per-sample data yield of 500 million bases retains sufficient power to detect a 1.5-fold increase in MF in our model. Thus, fewer animals per dose-group can be used with respect to the TGR models without losing power to detect an effect. Overall, our study provides strong evidence to support the use of DS in place of the plaque-based TGR assay for qualitative and quantitative chemical mutagenicity assessment with greater sensitivity, richer data and fewer animals.

### DS loci show Patterns of Mutation Sensitivity Consistent with TCR

There are many genomic and epigenomic features that have been linked to susceptibility to mutation induction including transcription (Monroe et al. 2022; Xia et al. 2020), sequence context (Hodgkinson and Eyre-Walker 2011), and chromatin state (Hodgkinson et al. 2012; Makova and Hardison 2015). The use of the Mouse Mutagenesis panel allowed us to observe differences in spontaneous and PRC-induced MF among the 20 targets. Specifically, we found that DS target loci in intergenic regions had a higher response to PRC mutagenesis than loci in genic regions, which is consistent with the protective effect of TCR. Transcribed regions of the genome are subject to increased rates of nucleotide excision repair (Nadkarni et al. 2016) that may contribute to the decreased MF in genic targets. Similar results were observed following BaP exposure in MutaMouse BM (LeBlanc et al. 2022). Remarkably, there was also a consistent pattern of relative sensitivity to mutation induction across targets between the two studies. In fact, the five targets with the highest MF_Min_ at the high PRC dose (chromosome 17, 14, 11, 8, and 16; all intergenic) were the same five targets with the highest MF_Min_ at the high BaP dose. These chemicals have different modes of action, which implies that there is an inherent feature of these loci that renders them more susceptible to mutagenesis than the others. Our results provide further support for the role of TCR in the variation of mutation among loci.

### Clonal Mutations and Target Responsiveness Underlie Discrepancy Between DS and LacZ

One of the main objectives of this study was to assess the concordance of DS with the gold standard TGR assay for *in vivo* mutagenesis testing. Initial analyses comparing the results of the *lacZ* assay with those of DS using MF_Min_ showed a significant positive correlation, but lower fold-increases in MF and a higher BMD for DS compared to *lacZ*. Similarly, an attenuated DS response compared to the Big Blue® plaque-based assay was observed in mouse BM analyzed 3 days following exposure to BaP (Valentine et al. 2020). These authors suggested that the difference might be attributed to unrepaired DNA adducts that were fixed into mutations during the *in vitro* portion of the TGR assay (i.e., *ex vivo* mutations). However, in this study, we had a 42-day waiting period between the last daily exposure and tissue collection. It is highly unlikely that DNA adducts could remain for such an extended period. A similar study using BaP also found the same difference following a 28-day sampling time (LeBlanc et al. 2022). Thus, *ex vivo* mutations are unlikely to contribute to the observed difference between DS and TGR assay. It is possible that applying the MF_Min_ approach to the DS data is too conservative and contributes to the observed difference between the two assay results.

The higher response measured by the *lacZ* assay may be the result of inherent differences between the two assays in how they quantify mutations. The *lacZ* assay scores mutants based on positive selection of plaques carrying mutant *lacZ* genes; thus, it cannot identify synonymous mutations, which do not alter the function of the *lacZ* gene, nor can it discount clonal expansion events unless paired with NGS sequencing. DS, on the other hand, identifies all types of mutations without bias and can easily filter out clonally expanded mutations using the MF_Min_ approach. A single mutation may produce large mutant frequencies following clonal expansion of the mutated cell (Heddle 1999). Due to the long interval between treatment and sampling time in our experiment, it is possible for the highly proliferative BM to accumulate small clonal populations of mutated cells. Indeed, we found that clonal mutations increased with PRC dose. When we included clonally expanded mutations in our calculations (using the MF_Max_ assumption), we observed that the concordance between DS and the *lacZ* assay improved significantly. DS fold-changes in several endogenous target loci became equivalent to fold changes observed with the *lacZ* gene. Furthermore, BMD confidence intervals generated using MF_Max_ overlapped with those for the *lacZ* BMD. Thus, it is possible that some of the discrepancies between DS and *lacZ* can be attributed to the exclusion of clonally expanded mutations when analyzing DS data. Whether clonal mutations should be included in the DS mutation counts (MF_Max_) or corrected for (MF_Min_) will require consideration of the context with which the assay is performed and should be a subject of further investigation by the expert community.

Differences in the responsiveness of the *lacZ* and DS assays to PRC may also be the result of assessing different genomic loci with differing sensitivities. Several studies have shown that exogeneous bacterial genes, such as the *lacZ* gene, tend to have higher mutant frequencies than endogenous genes. Previous work using Big Blue® mouse splenic tissue demonstrate that the *lacI* transgene exhibits higher spontaneous (Skopek et al. 1995) and BaP-induced (Skopek et al. 1996) mutant frequencies than the endogenous *Hprt* gene. Similarly, Valentine et al. (2020) found that the *cII* transgene has an elevated response to BaP mutagenesis compared to two expressed endogenous genes (*Ctnnb1, Polr1c*) using DS. We posit that the main reason for the seemingly attenuated response of DS compared to the *lacZ* assay is the result of the biologically higher susceptibility of the *lacZ* transgene to mutagenesis relative to many of the endogenous DS loci. The *lacZ* gene may exhibit increased susceptibility to mutation induction because it is a non-transcribed transgene, thus unprotected by TCR. Of the DS loci that had fold-increases in MF_Max_ rivalling that of the *lacZ* transgene, most were in intergenic regions. Future use of DS to directly sequence the *lacZ* gene would aid us in understanding how the differences between the two assays influence MF.

### DS Identifies Induction of Mutation Subtypes Not Feasible with TGR

A major benefit of DS is its ability to generate high-quality mutation spectra alongside its measurement of MF. Here, we showed that PRC induces mainly T:A>C:G, C:G>T:A, and T:A>A:T mutations. This is consistent with the known PRC mode of action as a monofunctional SN_1_-methylating agent (Fong et al. 1990; Fu et al. 2012; Gerson et al. 2018). The main mutagenic lesions produced by PRC include O_4_ and O_2_ – methylthymine leading to T:A>C:G and T:A>A:T mutations, respectively (Jenkins et al. 2005), and O_6_-methylguanine–a highly mutagenic adduct that results in C:G>T:A transitions (Fu et al. 2012). Previous work on the *lacZ* and *Pig-A* loci did not show significant increases in the proportions of C:G>T:A mutations following PRC exposure (Beal et al. 2020; Pletsa et al. 1997; Revollo et al. 2017). We believe our study is the first to report a significant increase in these mutations. This was likely possible because of the substantially larger amount of spectral data generated by DS than what was produced using laborious manual clone picking and sequencing with the other approaches.

Studies relying on TGR approaches characterized PRC mutation spectrum from only 20-120 mutations. In contrast, our study utilized a total of 3926 mutations, substantially improving the robustness of our analyses. Furthermore, spontaneous C:G>T:A mutations are extremely common at CpG sites as a result of spontaneous deamination of methylated cytosine to form thymine. We observed a dose-dependent proportional reduction in these mutations occurring at CpG sites, alongside an increase in the context of non-CpG cytosine bases, consistent with the formation of O_6_-methylguanine adducts by PRC. Previous studies with low mutation counts likely were underpowered to observe a significant increase in C:G>T:A mutations due to their abundance in the VC group. As a mutagenicity test, DS enables the detection of lower frequency changes in specific mutation types to facilitate the discovery of potential secondary mutagenic mechanisms or less prominent DNA lesions induced by the mutagen that would not be observed using traditional assays.

The trinucleotide mutation spectrum generated by DS following PRC exposure was linked to several COSMIC mutational signatures. The PRC mutation spectrum was most similar to SBS 21 and 26, which are both associated with defects in mismatch repair. Interestingly, the mutation spectra of similar acting alkylating agents (N-methyl-N-nitrosourea and N-ethyl-N-nitrosourea) within human pluripotent stem cells (Kucab et al. 2019) also showed high similarity with SBS 26 and SBS 12, suggesting that these SBS may be signatures of alkylation-induced chemicals. Additionally, we observed similarity with SBS 25, a signature observed in several Hodgkin’s cell line samples where the patients were treated with chemotherapy (Alexandrov et al. 2020). PRC was historically used to treat Hodgkin’s lymphoma; thus, it is possible this cancer signature is related to the exposure of these patients to PRC, or the similar acting methylating agent dacarbazine now used more commonly. These results support the utility of DS mutational signature analyses derived from mutagen-exposed rodents in identifying exposure-associated cancers in humans.

### DS Study Design Accommodates Reducing Animal Use and the Cost of Sequencing

The analyses of Piegorsch et al (1995) revealed that the main source of statistical variability observed in a TGR assay was significant variation between animals within a treatment group. These analyses provided the basis for the number of animals and plaques necessary to define a negative result with acceptable power (80%) for the TGR assay. OECD TG 488 requires a minimum of 5 animals per dose group for *in vivo* mutagenicity testing. In the current study, DS displayed much lower inter-animal variation than is typically observed using the *lacZ* assay. We note that the low sample-to-sample variability observed with DS is consistent with what other studies have found (LeBlanc et al. 2022). This prompted us to investigate whether the DS study design could be revised to accommodate a reduced number of animals. Our computations suggested that good power (80%) to detect a biologically significant (50%) increase in MF is attainable with as few as three animals and a minimum total depth of 500 million bases per animal. However, to observe more subtle increases in MF, such as that observed in the low PRC dose group, five animals per group is still required. Nevertheless, it appears that DS reduces intragroup variability, thus requiring fewer animals per dose group to detect an effect.

Studies with different variances would require different study designs. To ensure the robustness of our analyses to increased variance, we performed the same analyses with a hypothetical sample variance that was about twice as large as our observed variance (0.1 vs 0.06). We found that a minimum of three animals per group is still sufficient to achieve good power for a 50% increase in MF; however, the minimum number of bases sequenced per sample needs to be increased to 800 million to achieve this power. Overall, our analysis suggests that DS requires fewer animals per group than the TGR assay even when the variance is increased.

In the context of regulatory decision making, there remain many open questions on how to make a hazard call when targets within a panel vary in sensitivity to mutagenesis. For example, should a significant increase in mutations be observed in a single or a small subset of targets, but not with the overall panel, would this be sufficient for a positive result? Further work using DS to test weak and non-mutagenic chemicals across a broad panel of loci is needed to answer this question. Overall, the use of a specific custom panel of targets is an effective methodology to enable an assessment of the various genomic features that influence a chemical’s mutagenic effects.

## CONCLUSION

In summary, this study demonstrates the strong potential of DS for assessing *in vivo* mutagenicity and contributes key data towards efforts to understand optimal study design for its application as a regulatory test. With the inclusion of potentially clonally amplified mutations and the use of highly mutable targets, DS displays comparable fold-induction measurements to the gold-standard *lacZ* assay. In addition, DS provides high resolution mutation spectrum data that enable detailed investigations of mutagenic mechanisms and potential contributions to human carcinogenesis. Further advantages of DS with respect to the TGR assay include: (1) the potential to reduce the number of animals per group necessary to see a significant effect mitigating, in part, the costs associated with DNA sequencing; (2) possible integration with standard toxicity tests, thus negating the need for standalone mutagenicity with TGR animals, and further improving the efficiency and reducing costs and animal use for regulatory testing; and (3) application to any tissue from any species, including humans. Future work to establish DS as a mutagenicity test will require testing DS with different chemicals, including weak mutagenic and non-mutagenic chemicals, and examining response across different loci. Further studies are required to quantify relative mutational sensitivity associated with transcription status, chromatin state, and relationship to 3D genomic architecture across different cell types, tissues, and species to increase our understanding of the underlying biology of mutagenesis. Although there is still much work to be done to standardize DS, it shows great promise for regulatory mutagenicity assessment because it overcomes many of the current limitations faced by traditional assays and provides novel insights into chemical mechanisms that underlie cancer.

## Acknowledgments

The authors thank the extended team at TwinStrand Biosciences for their work and continuous bioinformatics and administrative support. Additionally, we thank the two Health Canada internal reviewers for helpful comments on the manuscript.

## TABLES & FIGURES

## SUPPLEMENTARY TABLES & FIGURES

**Supplementary Table 1.**
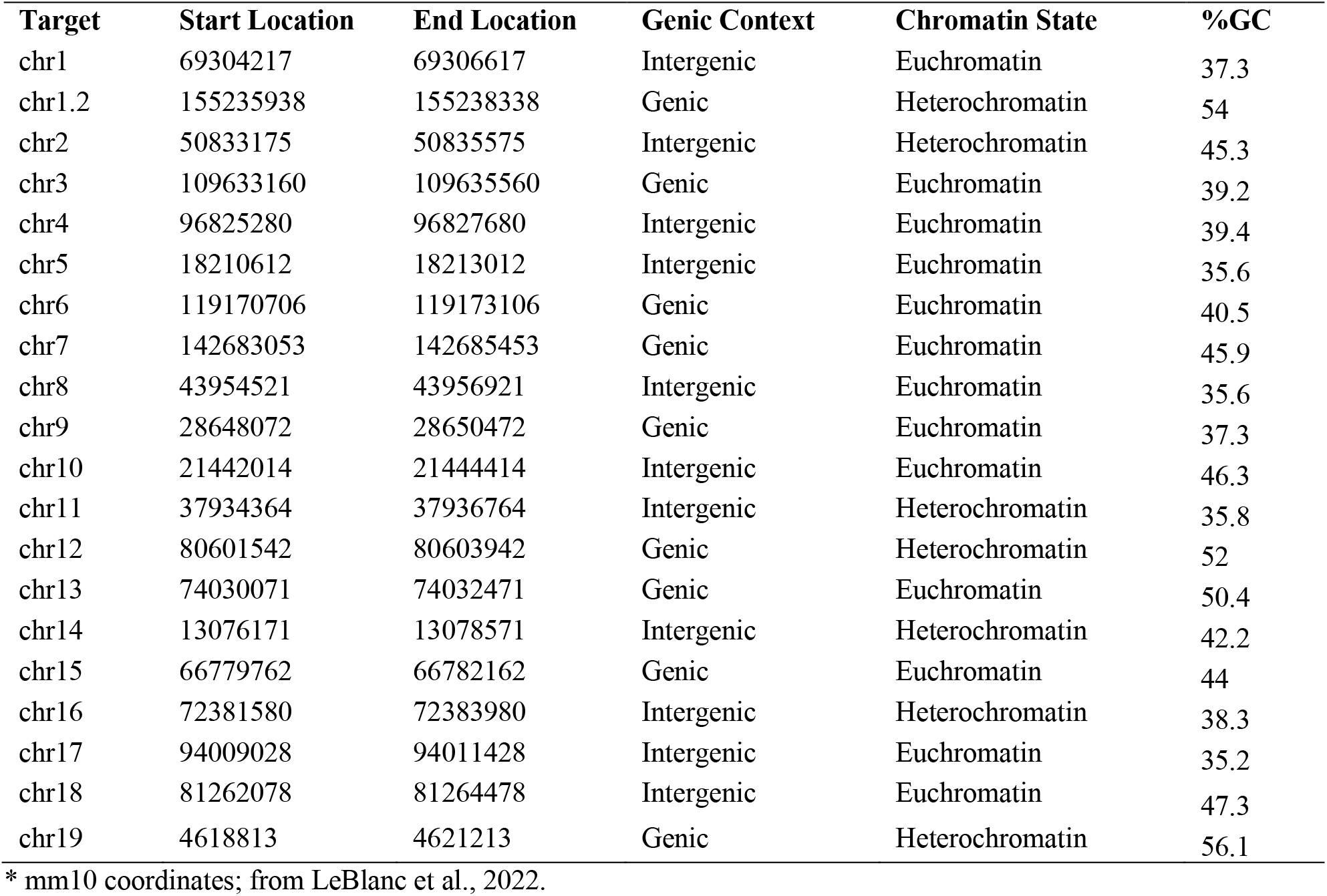
Mutagenesis panel targets and their chromosomal locations*

**Supplementary Figure 1.**
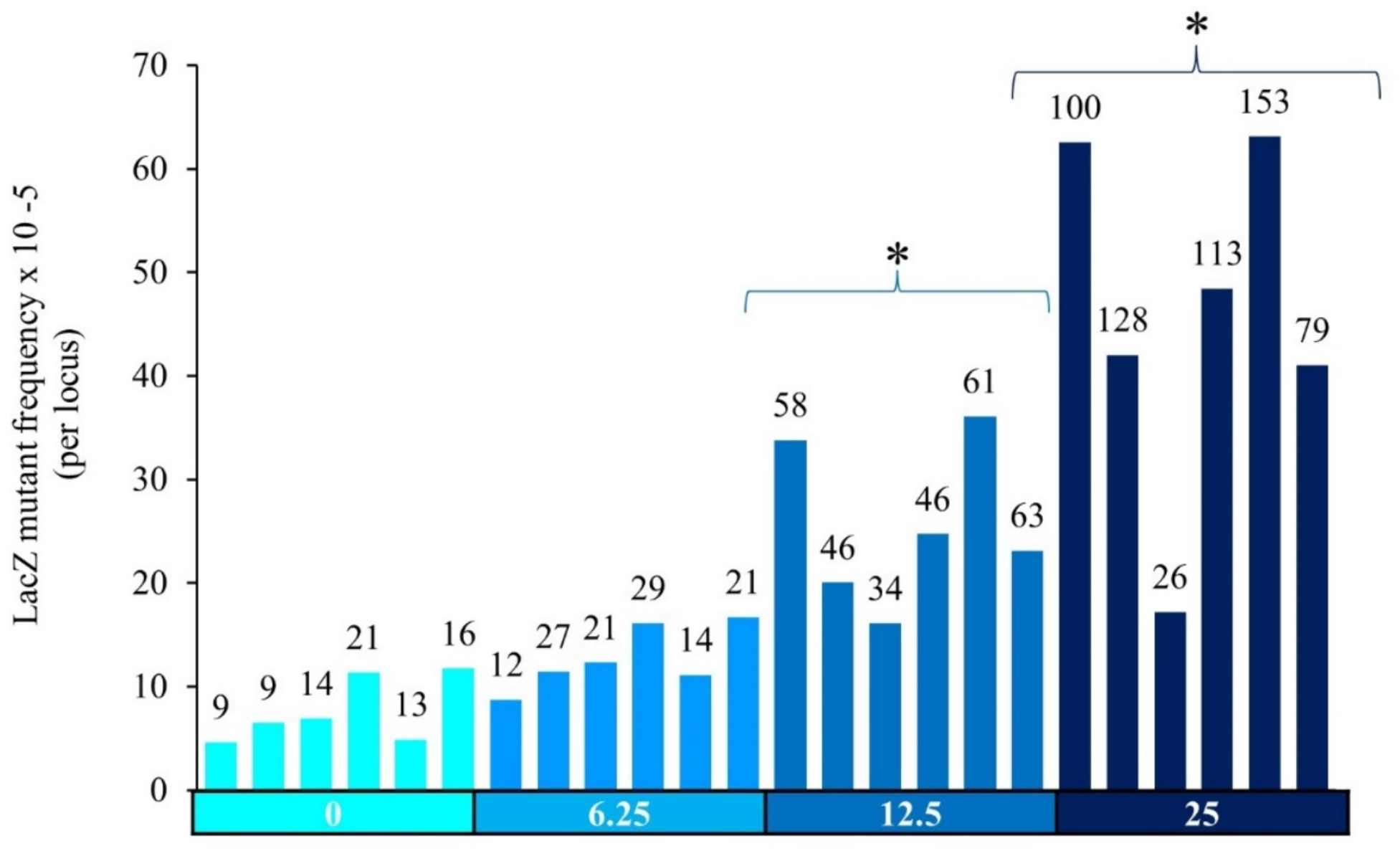
*LacZ* mutant frequencies in the bone marrow of MutaMouse males at various doses of PRC. Bars represent mutant frequency (mutations per locus) for each animal. Data labels indicate total number of mutant plaques counted for each animal. The X-axis indicates PRC dose group (mg/kg-bw/day). Asterisks indicate a significant difference relative the controls in the average mutant frequency across animals,.

**Supplementary Figure 2.**
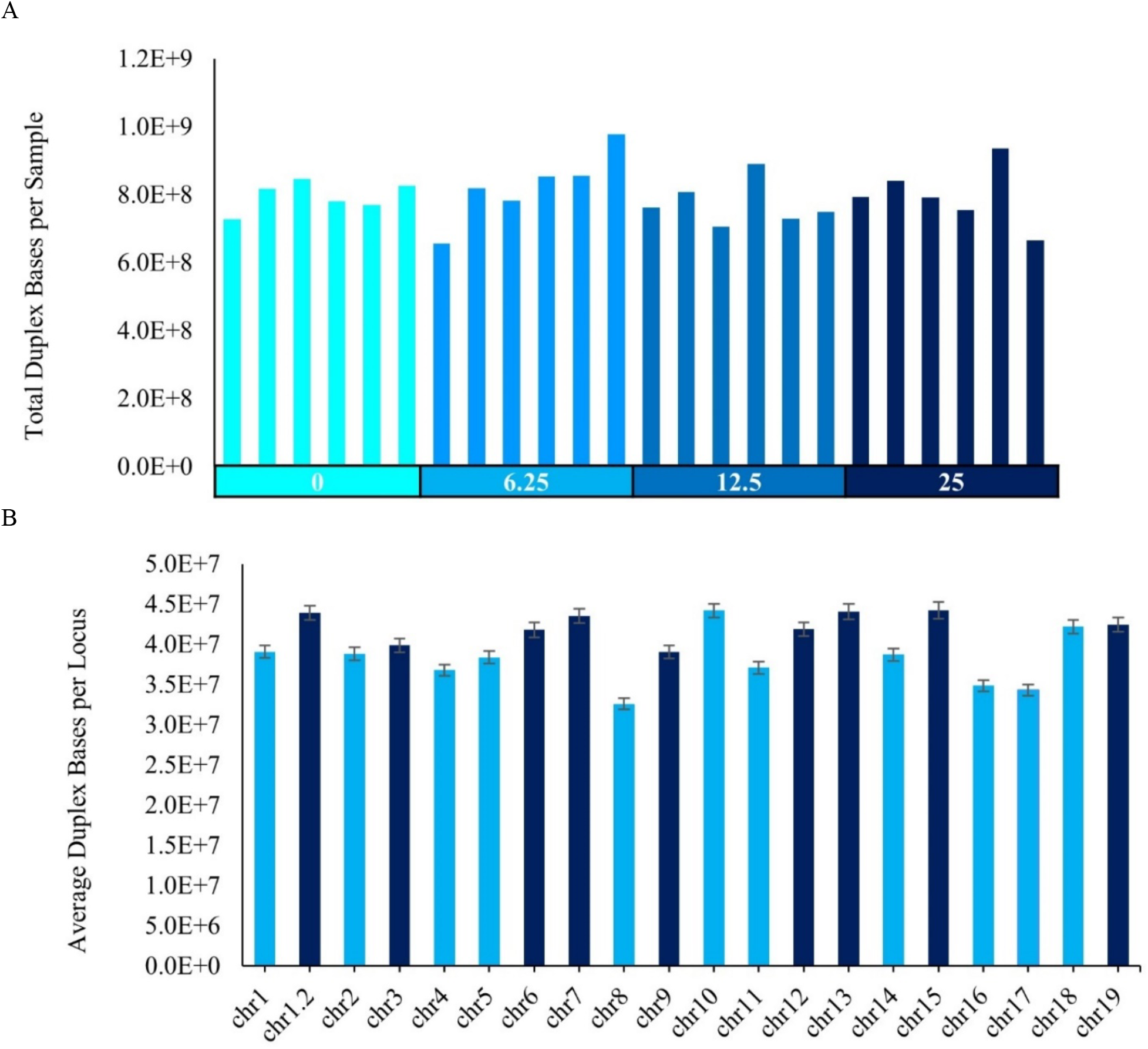
Duplex data yields. (A) Total duplex bases per individual animal. X-axis indicates dose group (mg/kg-bw/day PRC). (B) Average duplex bases per chromosome location per animal. Error bars represent standard error of the mean (SEM).

**Supplementary Figure 3.**
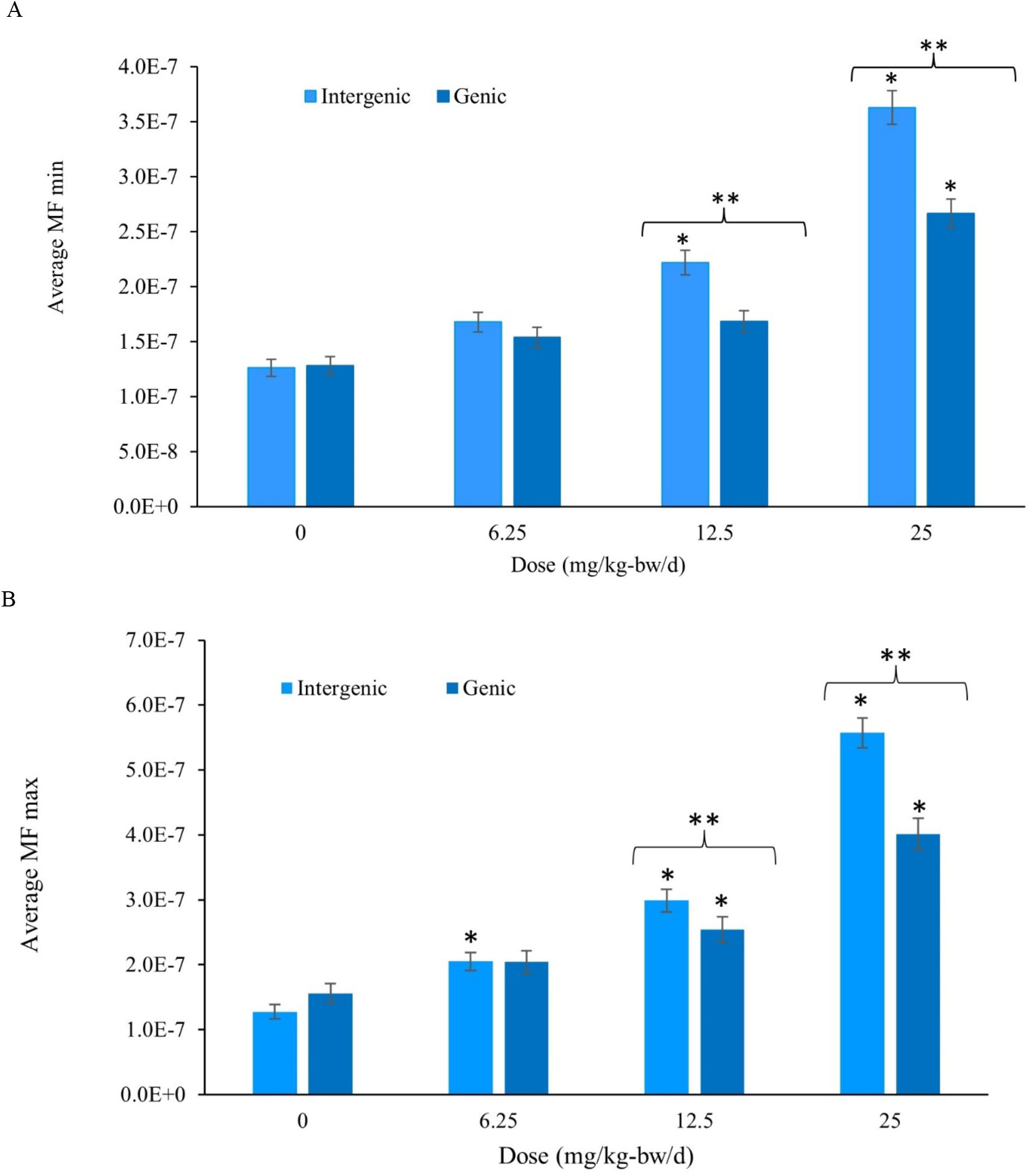
MF of intergenic targets compared to genic targets for various doses of PRC using A) MF_Min_ and B) MF_Max_. Data are represented as the mean ± SEM (mutations per bp). Asterisks indicate a significant difference in MF of dose groups relative to controls (GLM, p < 0.05). Double asterisks indicate a significant difference in MF between intergenic and genic targets within a dose group (GLMM, p < 0.05).

**Supplementary Figure 4.**
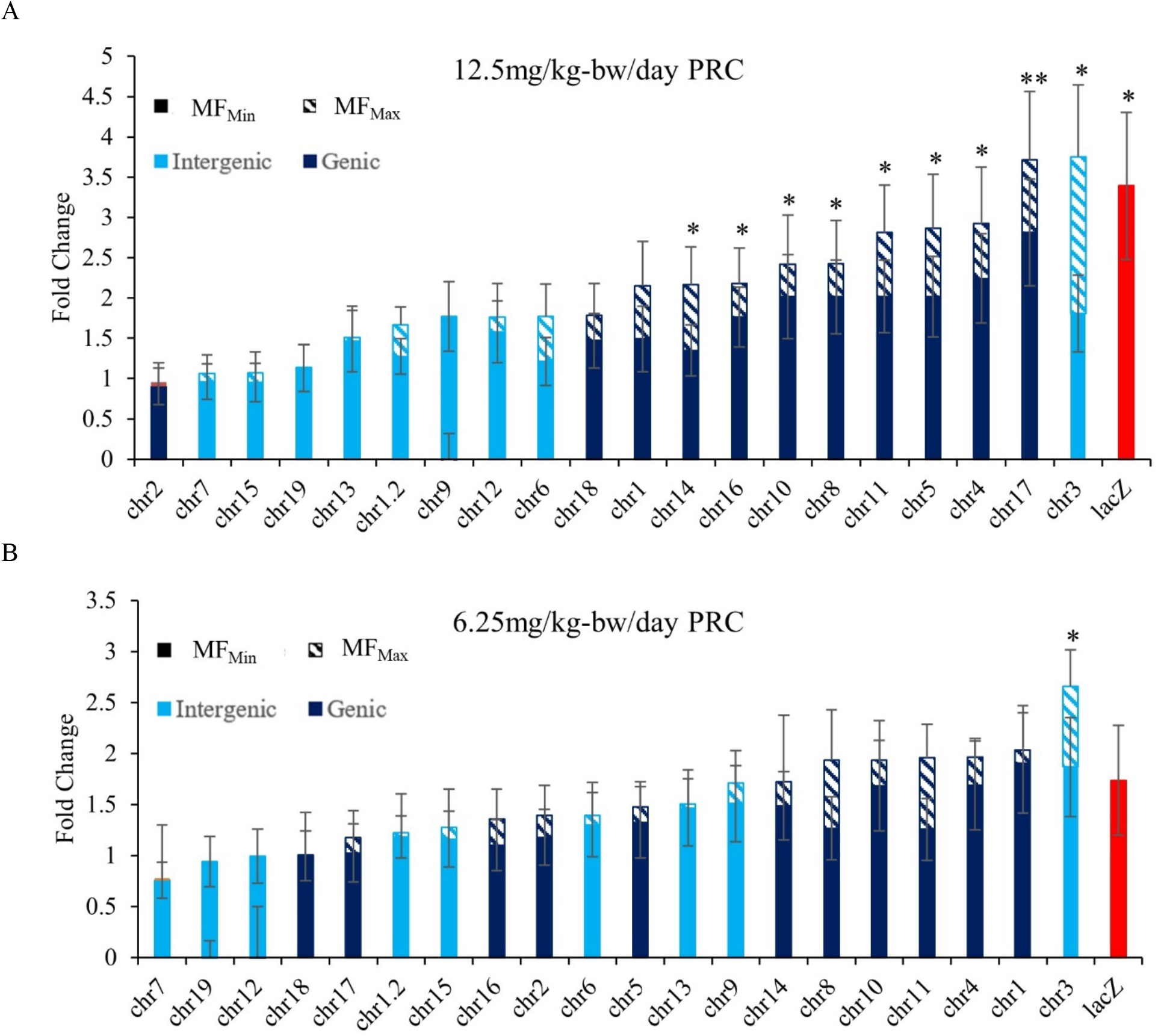
Fold change in MF between PRC VC (**A**) and middle-dose group (**B**) and low-dose group for the 20 DS targets as well as for the *lacZ* gene (red bar), estimated using a general linear mixed model. DS targets are listed along the X-axis. Errors bars are SEM. Asterisks indicate the fold change was a significant increase in MF_Max_ from VC. Double asterisks indicate significant increase for both MF_Max_ and MF_Min_.

**Supplementary Figure 5.**
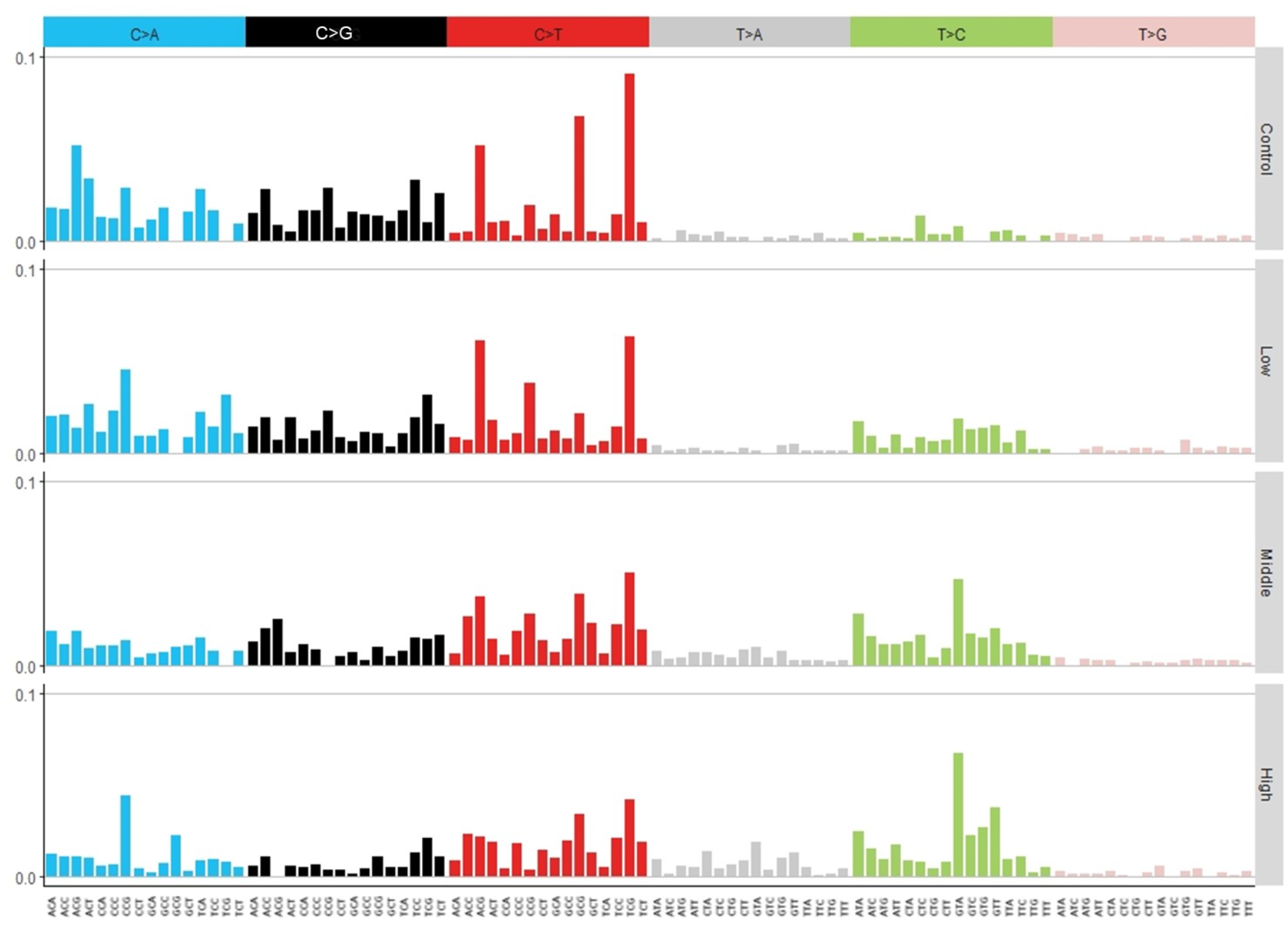
Mutation spectra with trinucleotide context of controls and PRC dose groups measured by Duplex Sequencing in the bone marrow of MutaMouse males. Dose groups are listed along the right: 6.25 mg/kg-bw/d (Low), 12.5 mg/kg-bw/d (Middle), 25 mg/kg-bw/d (High). The substitution subtype is listed along the top, the mutation including the two flanking nucleotides are listed along the bottom. Mutation subtypes are represented by the proportion of total mutations.

**Supplementary Figure 6.**
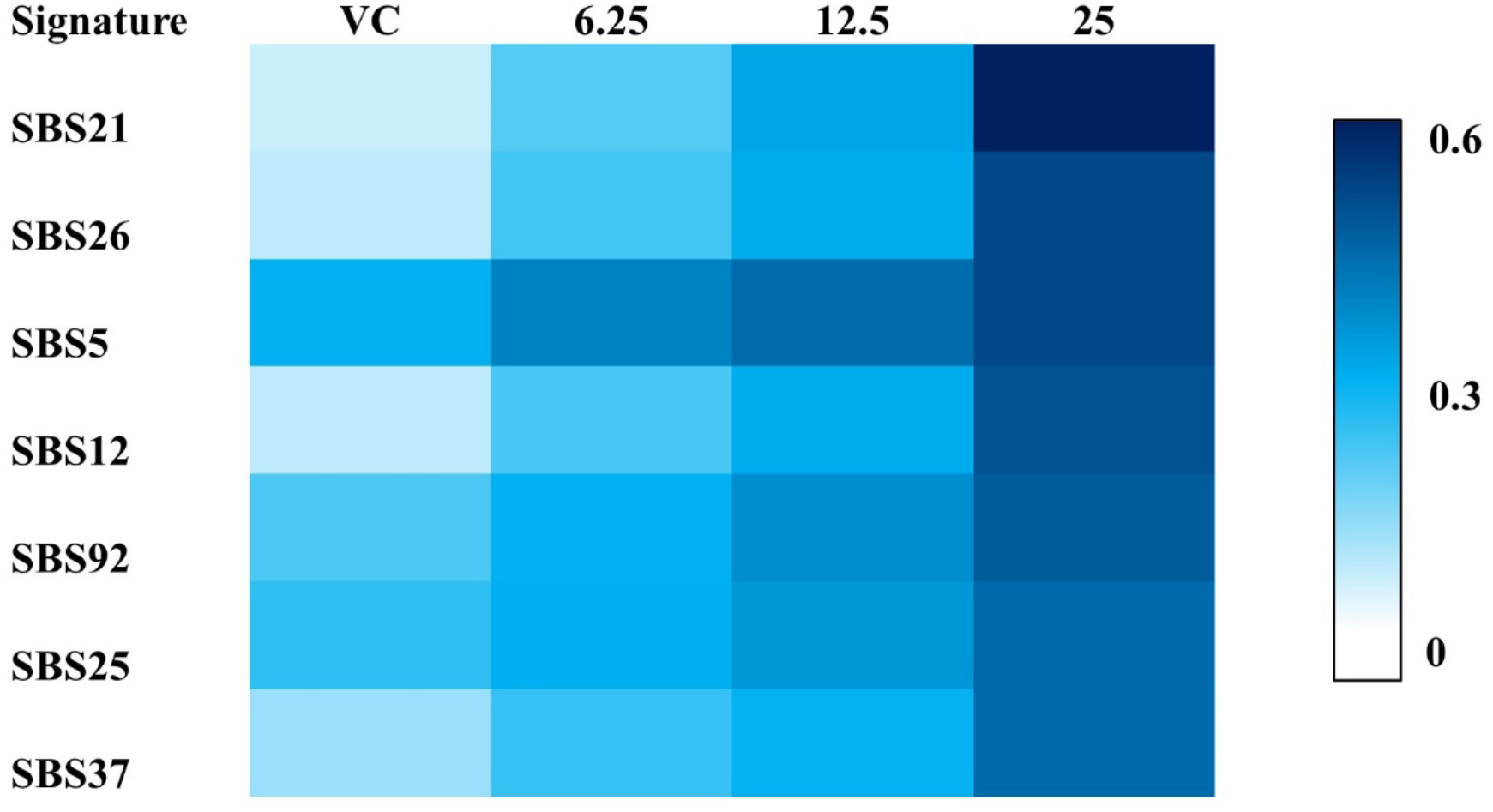
Cosine similarity of COSMIC SBS signatures to the trinucleotide mutation spectrum of each dose group. Data represents mean cosine value for the dose group.

## Notes

**Funding:** Health Canada’s Genomics Research and Development Initiative to FM. Additional funding support was provided from the Natural Sciences and Engineering Research Council of Canada (NSERC) and the Canada Research Chairs Program to CLY. AED received an NSERC Canada Graduate Scholarship and Ontario Student Assistance Program’s Ontario Graduate Scholarship to conduct this work.

**Conflicts of interest** P.V., F. Y. L., C.C.V., and J.J.S. are employees and equity holders at TwinStrand Biosciences, Inc. and are authors on one or more Duplex Sequencing-related patents.

### Competing Interest Statement

P.V., F. Y. L., C.C.V., and J.J.S. are employees and equity holders at TwinStrand Biosciences, Inc. and are authors on one or more Duplex Sequencing-related patents.

